# PSS: An enabling QTY server for designing water-soluble α-helical transmembrane proteins

**DOI:** 10.1101/724658

**Authors:** Fei Tao, Hongzhi Tang, Shuguang Zhang, Ping Xu

**Affiliations:** State Key Laboratory of Microbial Metabolism, Joint International Research Laboratory of Metabolic & Developmental Sciences, and School of Life Sciences & Biotechnology, Shanghai Jiao Tong University, Shanghai, 200240, People’s Republic of China; Center for Bits and Atoms, Massachusetts Institute of Technology, Cambridge, 02139, MA, USA

## Abstract

Membrane proteins, especially the α-helical ones such as G-protein coupled receptors (GPCRs), are considered extremely important owing to their significant biological roles. However, their expression and purification pose difficulties because of their poor solubility in water, which seriously impedes research progress in this field. Recently, QTY method, a revolutionary code-based protein engineering approach, was developed for the purpose of producing soluble transmembrane proteins. Here we describe a web server built for QTY design and certain analyses related to it (pss.sjtu.edu.cn). Typically, the Simple Design model is expected to take only 2-4 min, and the Library Design 2-5 h, of computer time, depending on target protein size and the number of transmembrane helices. Further, we describe a protocol for using the server with both Simple and Library Design modules. Protocols for experiments based on QTY design are also included. In summary, utilization of the web server, and associated protocols, will enable QTY-based protein-engineering to be implemented in a convenient, fast, accurate, and rational manner.

## INTRODUCTION

Membrane proteins, consisting of approximately 20%~30% genome-encoded proteins, are known to play vital roles in most of organisms(1). However, membrane proteins are still heavily under-represented in protein data banks owing to serious difficulties encountered in the expression and purification of these proteins because of their poor solubility(2). The α-helical transmembrane (TM) proteins comprise the major category of membrane proteins. It is estimated that around 27% of all human proteins are α-helical membrane proteins(3), among which, the GPCRs are most interesting(4,5). In our recently published study, which used GPCRs as model proteins, a code-based approach, named the QTY method, was successfully developed for solubilizing TM proteins. It is considered a revolutionary new method for making a membrane protein water-soluble(6). The method was tested by making several GPCRs, including CCR5, CXCR4, CCR10 and CXCR7, water-soluble. The results indicated that all artificial GPCR variants are water-soluble and able to maintain ligand-binding activity by slightly changing K_m_ values. This method may be extensively used to modify other membrane proteins containing TM helices, including transporters and ion channels.

Although the QTY strategy is straightforward in comparison to previous methods, doing QTY design manually is still time-consuming and tedious. The library design, especially, is virtually impractical to complete without computing power. Moreover, QTY related bioinformatics analyses are extremely tedious, as 7 different software programs are involved and the resulting output needs to be integrated. Therefore, PSS, a web-based server, was developed to facilitate the QTY method by avoiding the tedious analyses involved. The web-based server was developed to provide a graphical and friendly interface to a broader group of users. Software programs for QTY-related analysis were incorporated into the server to prepare elaborate analysis reports for QTY design. Using PSS for typical QTY Simple Design, a standard analysis report can be completed within 4 minutes and only requires very simple operations such as copy and pasting. PSS is expected to be widely used by scientists endeavoring in the research about α-helical proteins.

## METHODS AND IMPLEMENTATION

The workflow and architecture of PSS server are illustrated in Figure 1. The following subsections describe the main features of each step/part in detail.

**Figure 1.**
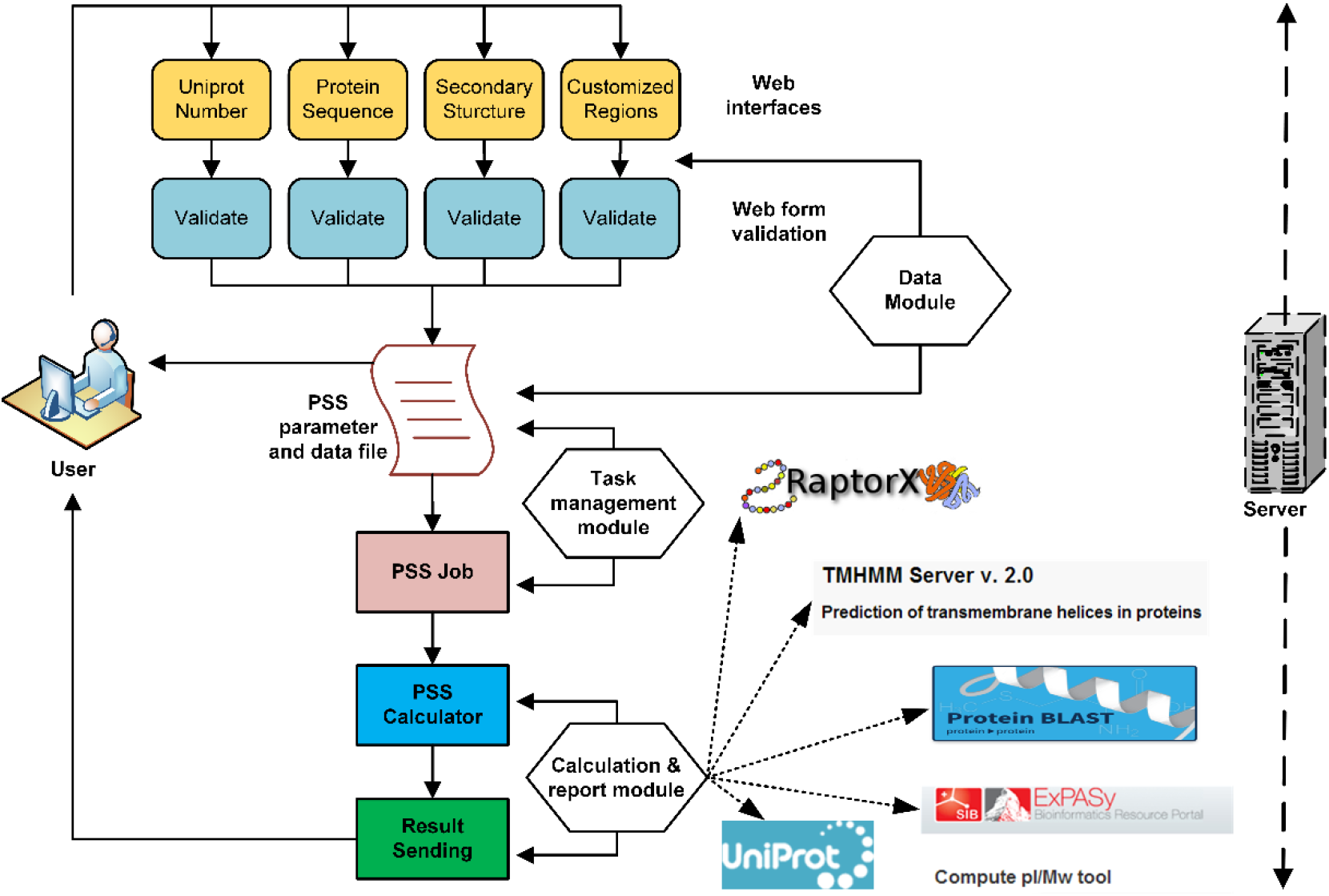
Flowchart of PSS server. The web server consists of 4 parts; namely web interface, input data managing module (Data Module), task managing module (Task management module), and the core calculation module (Calculation and report module). In the core calculation module, different software were introduced to achieve versatile function. ExPASy is presented as an example of the software used for the prediction of pI and M_W_ values. The software ProPAS and Protter server are not shown although they were also used by PSS. The hydrophobicity values were calculated using ProPAS.

### Input

Two types of information are needed for QTY design. These are the protein sequence and its corresponding TM region information (Fig. 2). By default, a user input must include a UniProt number, which is the unique ID of a protein record in the UniProt database. Then, the server will retrieve that protein sequence from the UniProt database (https://www.uniprot.org/), and use it for QTY substitutions. If multiple sequences are represented by a single UniProt number, only the canonical sequence will be used for the design. Alternatively, a user can also input a protein sequence by simply typing/pasting that protein sequence in the provided text field (Supplementary File 1).

**Figure 2.**
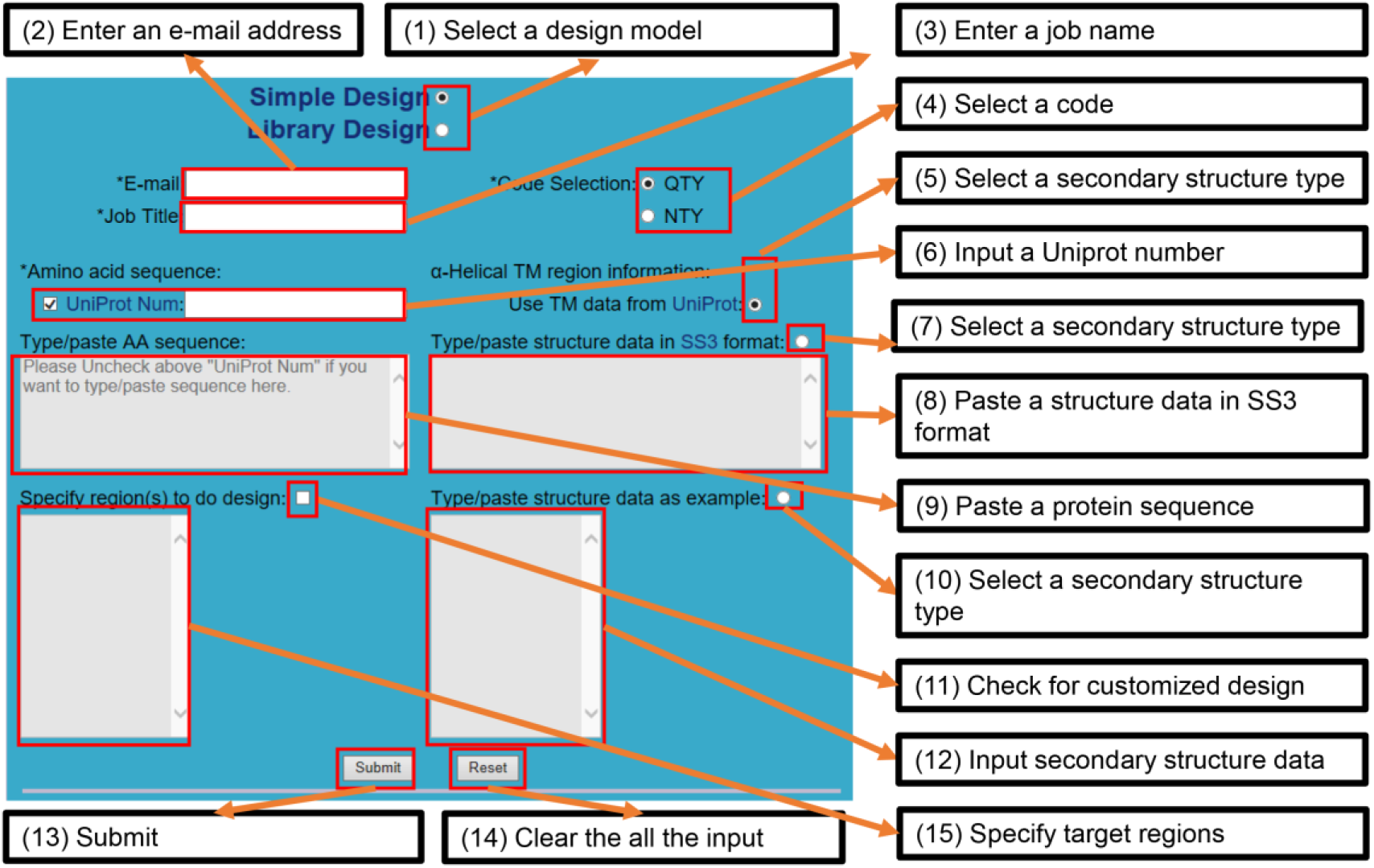
Screenshot of the PSS Design page. The major parts of the web interface are shown. It includes 6 sections, which contains a total of 15 elements such as text fields, and check boxes. The asterisk is used to label required items. UniProt is the name of the protein database (www.uniprot.org). The ‘/’ between ‘type’ and ‘paste’ means ‘or’. The text fields in gray are disabled by default.

With regard to TM region information, the design becomes more complicated. Generally, it is difficult to predict the type of data which is more accurate. Therefore, entering TM region information in different way is allowed. If the UniProt database is used as the data source, the TM region information from the UniProt database will be automatically selected for QTY design (Fig. 2). Alternatively, there are 2 other exclusive manual ways, which are; (i) pasting a SS3 format string, and (ii) manually indicating the start and end positions of all TM regions (Supplementary File 1). For typing/pasting a sequence as the input, there are 3 ways of getting the TM region information. Apart from the 2 manual ways stated above, the server may also obtain TM information by performing TM region prediction using TMHMM V2.0, developed by Anders Krogh and rated as the best way to predict TM helices, when no external TM region information is available(1,7,8).

Considering that partial modification of a protein may be required by users at some point, PSS also allows the manual selection of specific TM regions to perform a QTY design. For this, a user is only required to input the start and end positions of each selected protein fragment (Fig. 2). In such a situation, the server will only perform QTY substitutions in the TM region(s) within the indicated fragment(s).

### Simple-Design

This module, developed for classical QTY design, was named Simple Design. It conducts a direct and complete substitution of amino acid residues in a protein with a selected QTY code (Fig. 3). After obtaining the input sequence and TM region information, the server will substitute all changeable residues in the TM regions and output a sequence for the designed protein. By default, QTY code is selected, but it can be changed if a user desires to use NTY code. The difference between the 2 codes is that the amino acid N instead of Q is used for substitution of amino acid L with NTY code(6).

**Figure 3.**
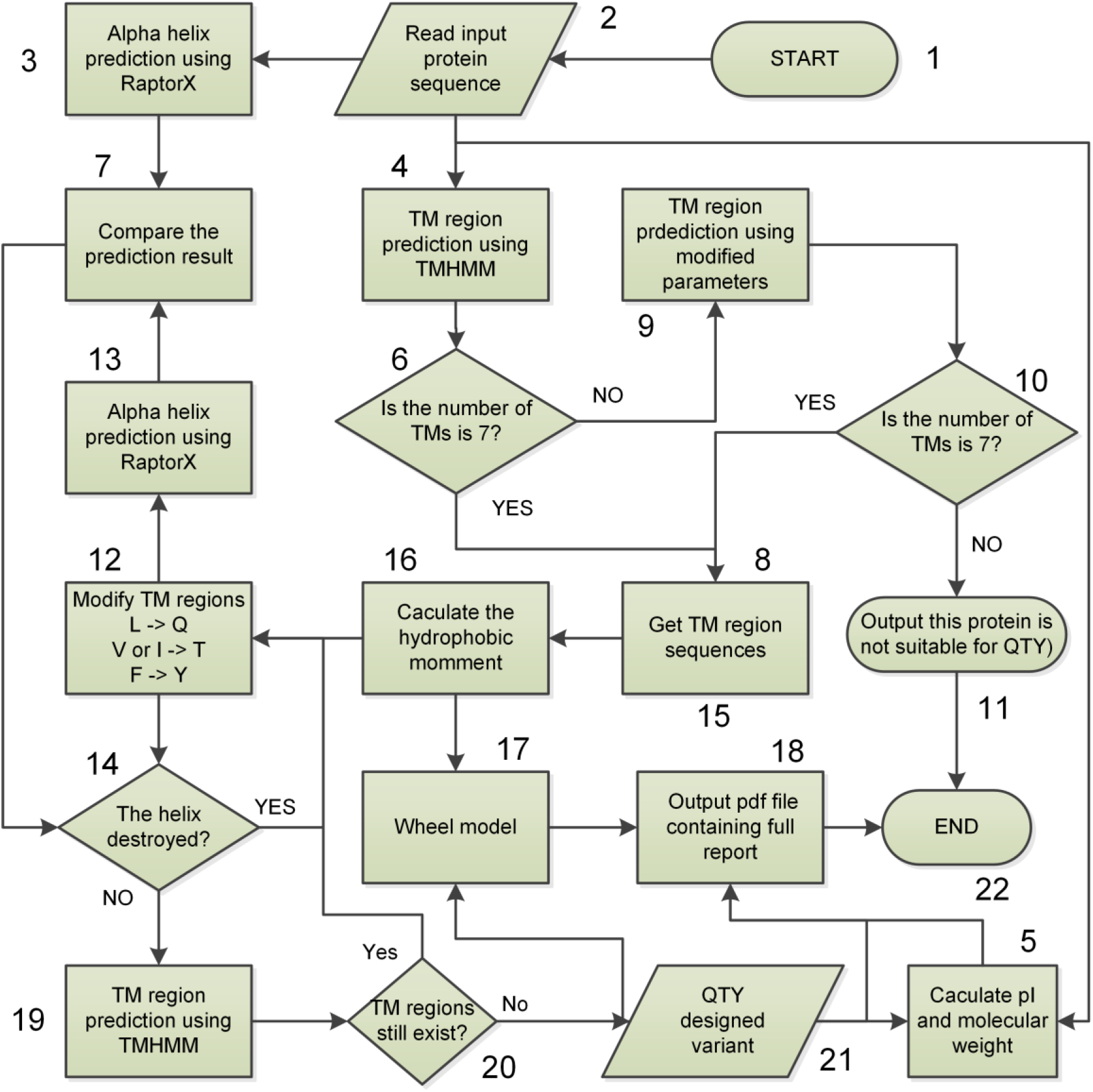
Flow chart for QTY protein design. The work flow for 7-TM GPCR design was selected as an example to show the computing process of the Simple Design in case only sequences input by the user are available. This is the typical work model of PSS. The process can be divided into 22 steps including the input and output. All steps were numbered to facilitate description. The label ‘->’ means ‘to’.

After the input, QTY substitution can be completed in seconds, because there is no time-consuming calculation step. For the Simple Design, the main cost in time is due to the comparisons that are performed to display details of the design in the output report. There are 5 comparisons in a standard Simple Design report (Supplementary File 2). The first one comprises general characteristics including calculated molecular weight (M_W_), isoelectric point (pI), and hydrophobicity (H_Y_). These calculations are made using ProPAS software, and ExPASy server, developed previously(9,10). Comparisons of 15 well-elucidated human GPCRs and their QTY-designed variants are shown (Table 1). All comparison data were directly copied from standard Simple Design reports. The second comparison is based on TM region prediction, achieved using a standalone version of the TMHMM V2.0 software(1,8). Detailed sequence alignments of a QTY designed protein and its origin was done using a Perl script, where a Perl SVG module(11) was used to draw the α-helix schematic. An interactive protein feature visualization software, Protter(12), was also used to show the difference between the original proteins and the designed ones. This software is also able to predict TM regions, while it uses a serpentine-like map to show the secondary structure pattern of a protein. The membrane localization of a protein is also predicted and shown in the output map. Moreover, a detailed comparison of each helix will also be shown in the report. RaptorX-Property (previous RaptorX-SS8)(13) software, ranked first in secondary structure prediction(14), was incorporated to perform above comparison. Sequence alignments of TM regions and alignments of α-helix prediction results are shown in the last section of the report. In this section, the helical comparisons are shown using helical wheels of all TM regions. A modified helical wheel drawing software was used to prepare the wheel figure (http://rzlab.ucr.edu/scripts/wheel/wheel.cgi). All comparisons are integrated and presented in a PDF format report file that is to be sent to the user’s email address (Supplementary File 2).

**Table 1.** Selected human GPCRs and their QTY-designed variants.

| UniProt<sup>a</sup> | Wild Type (WT) |  |  | QTY-designed (MT) |  |  | R<sub>ACT</sub><sup>e</sup> | R<sub>SC</sub><sup>f</sup> |
| --- | --- | --- | --- | --- | --- | --- | --- | --- |
|  | M<sub>w</sub><sup>b</sup> | pI<sup>c</sup> | H<sub>y</sub><sup>d</sup> | M<sub>w</sub> | pI | H<sub>y</sub> |  |  |
| P41595 | 54.30 | 9.22 | 0.2672 | 54.76 | 9.10 | -0.7305 | 0.5484 | 0.0291 |
| P29274 | 44.71 | 8.33 | 0.4607 | 45.15 | 8.29 | -0.7072 | 0.5060 | 0.1068 |
| P07550 | 46.46 | 6.59 | 0.1700 | 46.65 | 6.59 | -0.8764 | 0.4731 | 0.0630 |
| P51681 | 40.52 | 9.20 | 0.5466 | 41.05 | 9.07 | -0.9218 | 0.5576 | 0.0483 |
| P61073 | 39.75 | 8.46 | 0.3994 | 40.05 | 8.40 | -0.9113 | 0.5608 | 0.0568 |
| P35462 | 44.22 | 9.20 | 0.3190 | 44.71 | 9.12 | -0.7351 | 0.4902 | 0.0750 |
| P41143 | 40.37 | 9.20 | 0.5384 | 40.75 | 9.11 | -0.6878 | 0.4629 | 0.0538 |
| P47871 | 51.26 | 9.01 | 0.0785 | 51.93 | 8.90 | -0.9388 | 0.5094 | 0.0929 |
| P35367 | 55.78 | 9.33 | -0.0859 | 56.20 | 9.24 | -0.9675 | 0.5067 | 0.0924 |
| P41145 | 42.65 | 7.92 | 0.4816 | 42.85 | 7.88 | -0.7465 | 0.5241 | 0.0500 |
| P41146 | 40.69 | 8.73 | 0.6511 | 41.14 | 8.67 | -0.4883 | 0.4720 | 0.0243 |
| Q9H244 | 39.44 | 9.59 | 0.3512 | 39.85 | 9.38 | -0.9786 | 0.5490 | 0.0263 |
| P21453 | 42.81 | 9.58 | 0.4408 | 43.38 | 9.44 | -0.9009 | 0.5506 | 0.0314 |
| P28222 | 43.57 | 8.95 | 0.3421 | 43.99 | 8.88 | -0.8378 | 0.4938 | 0.0590 |
| Q99835 | 83.68 | 8.70 | -0.0599 | 84.17 | 8.66 | -0.5953 | 0.5102 | 0.1329 |

<sup>a</sup>All the 15 selected GPCRs have been well investigated in previous research, and their related structure data can be retrieved in PDB. <sup>bcd</sup>The M<sub>w</sub>, pI, and H<sub>y</sub> indicate molecular weight in kDa, isoelectric point, and hydrophobicity respectively, which were calculated using ProPAS(9).
<sup>e</sup>The R<sub>ACT</sub> is for estimating the changing rate of amino acid sequence in TM regions, which was calculated through dividing the numbers of changed amino acids by summarized length of all the TM regions. <sup>f</sup>The R<sub>SC</sub> is for showing the changing rate of secondary structure, which was calculated through dividing the number of positions related to structure-changing by the whole length of the protein sequence.

### Library-Design

In the Simple Design module, all changeable amino acid residues within TM region(s) are substituted. Therefore, probability of designing a soluble protein is high. However, there is also high probability that the designed protein may loss its original functions. The rule that applies is: as more AA residues are changed, the possibility that the designed protein is soluble will increase, but the possibility that the protein is non-functional will also increase. Therefore, only the minimum number of amino acid residues that need to be changed to make the TM soluble must be changed, so that the key amino acid residues needed to maintain protein functions are retained untouched. This emphasize the need to find a balance between solubility and protein structure integrity.

To accomplish the above, a work flow program (Fig. 4) was developed, based on the domain shuffling principle (Fig. 5), to randomly change TM residues and screen all variants using a high-throughput procedure in silicon. There are approximately 85 changeable amino acid residues for a typical GPCR. Therefore, theoretically, there may be 2^85 random variants of a typical GPCR protein, which makes both library construction and screening impractical. Thus, a compromise approach using partially random method was developed for our server. In this approach, TM region prediction software TMHMM(1) and secondary structure prediction software RaptorX(15) were combined to rule out most insoluble or non-functional variants in silicon. TMHMM was used to determine if a modified α-helix is water-soluble, and RaptorX is used for determine if the modified TM region could still form α-helical structure. A scoring method was introduced to evaluate the balance. An example of the design of a typical 7-TM GPCR protein is shown (Fig.5). As shown, there are 8 different designed variants for each TM region and The NTM (Non-TransMembrane) region is the same as that of the original protein. There are 64 fragments of amino acid sequences in total. Next, the different domains are randomly integrated to form different full-length variants. Calculations indicate approximately 2 million (the 7th power of 8) QTY variants for each 7-TM GPCR library. In silicon filtering step in the Library Design is highly time-consuming and typically takes approximately 2 h for a single library design of a GPCR containing 7 TM regions.

**Figure 4.**
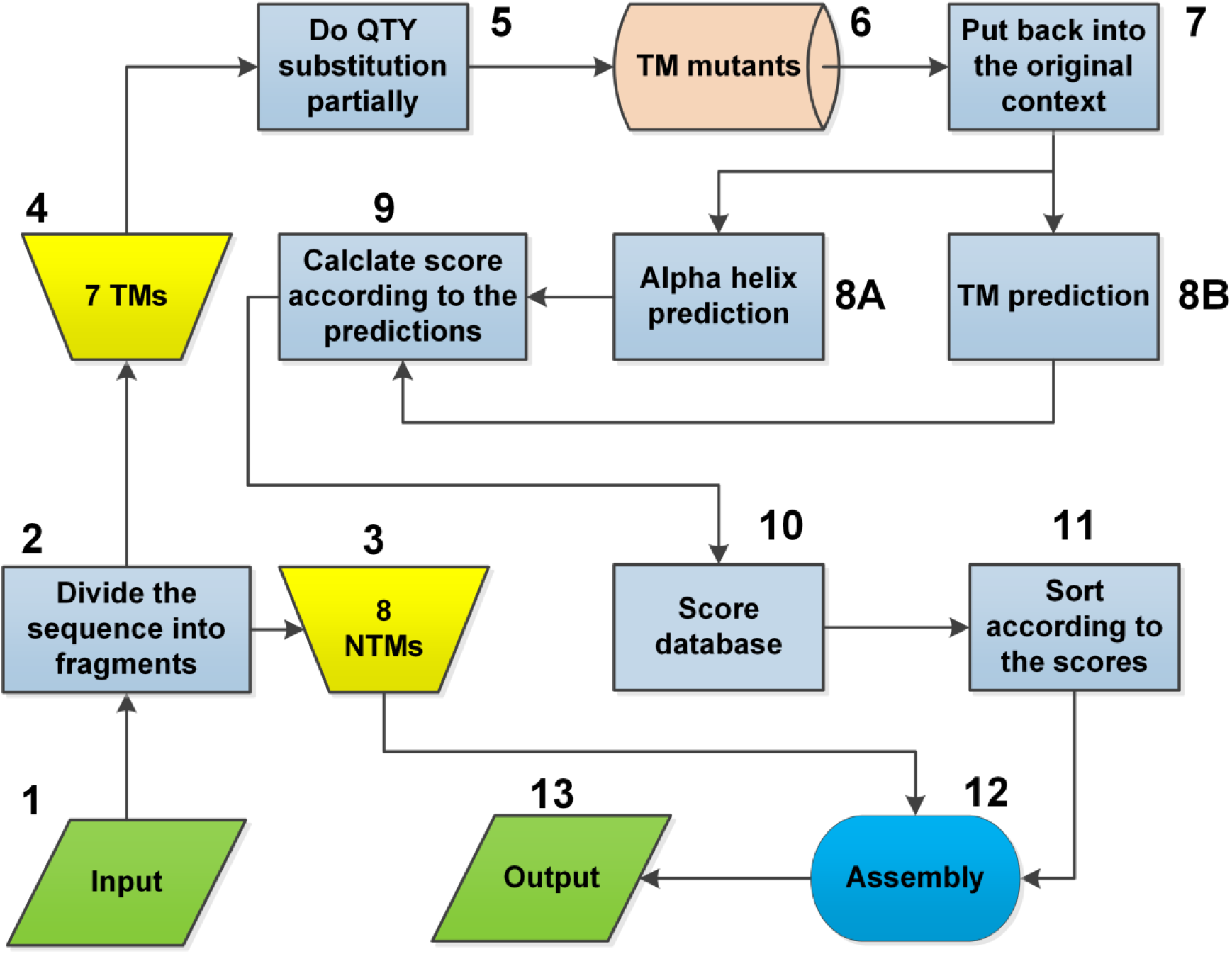
Flow chart for QTY-Y2H library design. The work flow of library design is shown using 7-TM protein as an example. NTMs indicate non-transmembrane regions in a protein. The number represents the order of design steps. There are 13 steps in total, including the input and output steps. TMs indicate transmembrane helices.

**Figure 5.**
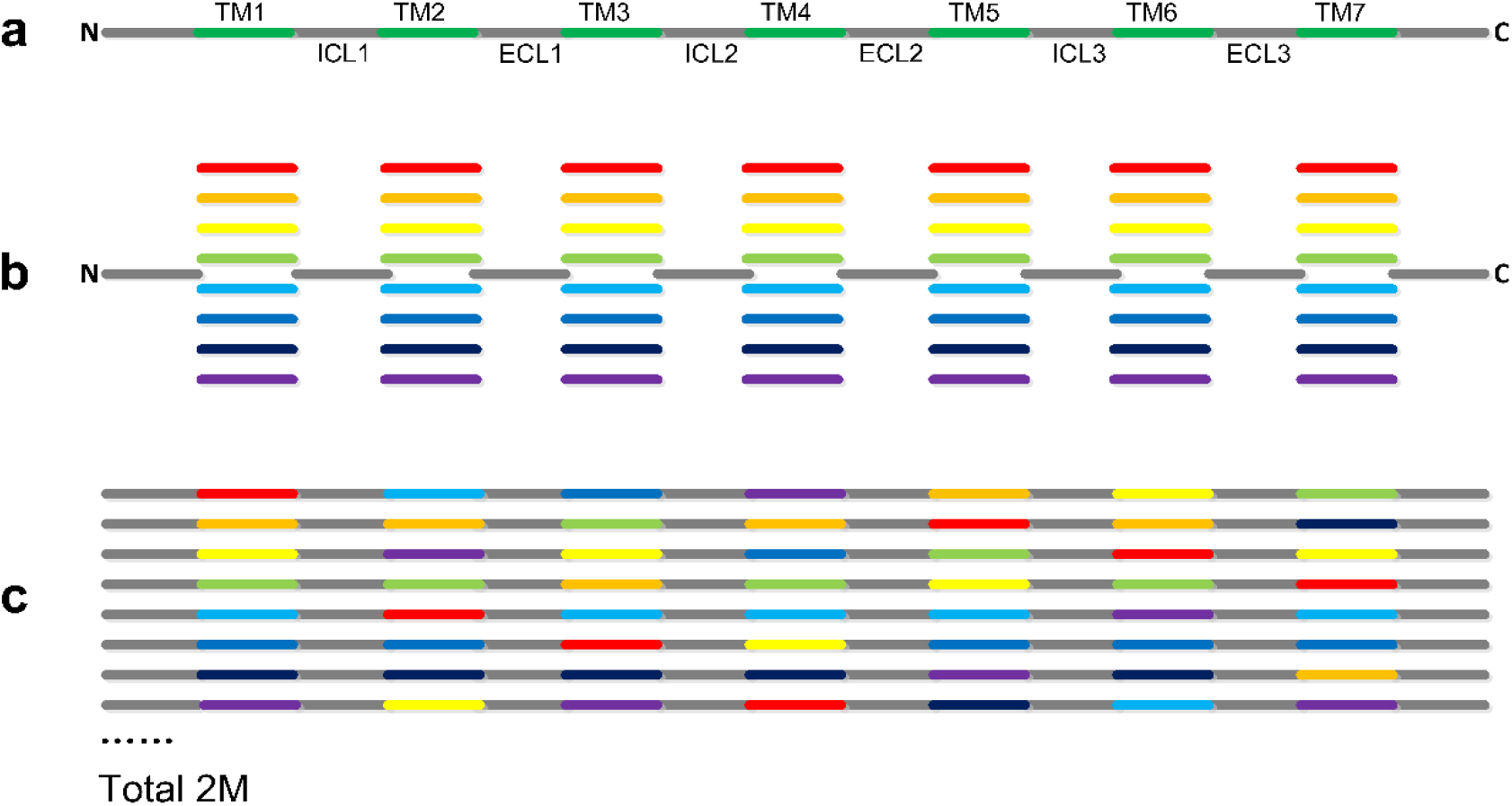
Principle of directed library construction. The design was divided into 3 steps; a, b and c. (a) A typical 7-TM protein was divided into 15 fragments including 7 TMs (TM1-7), 3 intra-cellular loops (ICL1-3), 3 extra-cellular loops (ECL1-3), 1 N-terminus and 1 C-terminus. (b) Design 8 variants for each TM. (c) All fragments are assembled together, and each variant is used randomly. ‘Total 2M’ indicates the total number of all combinations, which is over 2 million (8^7).

Considering the time-consuming nature of the library design, we deposited 126 previously designed libraries of GPCRs in the server’s database. Some selected proteins well screened for their physiological roles in health and disease, are shown (Table 2). The complete list of deposited libraries is included in the supplementary file (Supplementary File 5). These are all the known human GPCRs that possess protein/peptide ligands. The library designed using these proteins may be screened using the yeast two hybrid (Y2H) method(16), which is suitable for GPCRs with protein/peptide ligands that have been successfully used in our previous work(6). By using the UniProt number as input and using default parameters, the deposited library designs can be immediately extracted, and sent to the user’s email address. Alternatively, the server will run the of Library Design pipeline from scratch, and forward an email report to the user upon completion of the job.

**Table 2.** Selected 7-TM GPCRs suitable for Y2H library design.

| Receptor Name | UniProt Ref (R)<sup>a</sup> | Size | Ligand Name | Ligand Size | UniProt Ref (L)<sup>b</sup> | Family Name |
| --- | --- | --- | --- | --- | --- | --- |
| CXCR4 | P61073 | 352 | SDF-1α | 65 | P48061 | Chemokine receptors |
| PAR2 | P55085 | 397 | Serine proteases | >100 | / | Proteinase-activated receptors |
| V3A receptor | P30518 | 371 | Arg-vasopressin | 9 | P01185 | Vasopressin and oxytocin receptors |
| CCR5 | P51681 | 352 | CCL11,14, 16, 2, 3, 4, 5, 7, 8 | 74 | P51671 | Chemokine receptors |
| ETB receptor | P24530 | 442 | Endothelin-1, 2, 3 | 21 | P05305; P20800; P14138 | Endothelin receptors |
| MC1 receptor | Q01726 | 317 | α-MSH | 13 | P01189 | Melanocortin receptors |
| ACKR3 | P25106 | 362 | CXCL11, SDF-1α | 70 | O14625; P48061 | Chemokine receptors |
| PTH1 receptor | Q03431 | 593 | PTH; PTHrP | 84; 141 | P01270; P12272; P12272 | Parathyroid hormone receptors |
| Kisspeptin receptor | Q969F8 | 398 | Kisspeptin-10, 13, 14, 52, 54 | 10-54 | Q15726 | Kisspeptin receptor |
| CT receptor | P30988 | 508 | Calcitonin; amylin | 32; 37 | P01258; P10997 | Calcitonin receptors |
| CCR2 | P41597 | 374 | CCL2 | 76 | P13500 | Chemokine receptors |
| GnRH receptor | P30968 | 328 | GnRH I, GnRH II | 10 | P01148; O43555 | Gonadotrophin-releasing hormone receptors |

<sup>ab</sup>The 'R' in braces stand for receptor, and 'L' for ligand.

There are 2 TXT format files in the standard report of the Library Design (Supplementary File 3). In one file, the sequences are separated into multiple lines for ease of printing and reading using a PC. In the other file, each sequence is presented as single long line, for the convenience of copying and editing using a computer. In our study, single-line files were sent to the company for their service. Notepad++ is strongly recommended for opening the output files. Any other software compatible with TXT file editing may also be used.

### DNA synthesis

A user of the PSS will receive an output of protein sequence(s). Next, it is usually required to design the corresponding DNA sequence(s), which can be used in subsequent protein expression and characterization experiments. In the Simple Design module, only one sequence is put out by the server. Therefore, a user is only required to perform a reverse-translation for the following cloning and expression, with the translation codon usage bias in consideration. This may be performed with bioinformatics tools such as JCat(17) and OPTIMIZER(18), which have been repeatedly and successfully applied in our lab. This work may also be achieved through a commercial service, which is much convenient for most users.

For synthesis of the library, a user must seek help from a professional company experienced in DNA synthesis. Usually, most companies do not have a well-established method for library synthesis. In our previous study, we aided 2 companies to establish the library synthesis approach. The service of these 2 companies, GenScript (Nanjing, China) and Ginkgo Bioworks (Boston, USA, previous Gen9), are available worldwide. In brief, the synthesis process is as follows: Each of the 64 fragments of AA sequences is reverse translated to the corresponding DNA sequences; next, overlap sequences are added to the ends of each DNA fragment for assembly; then, the DNA is synthesized and assembled using PCR. The final PCR product will be a mixture containing approximately 2 million variants (8^7). This library can then be used in the follow up high-throughput screening such as Y2H.

### Protein expression

Theoretically, any strategy or host s for protein expression may be used for expressing the designed protein. Therefore, users are encouraged to try different methods in performing protein expressing experiments. Here we will only be providing some advices for users based on our previous experiments. For expression of QTY designed human GPCRs, we have used *E. coli*, and insect cells as hosts. Insect cells appear to be highly suitable for expression, while using *E. coli* always causes difficulties related to inclusion-body issues. Other hosts system, such as the yeasts may also be suitable because Y2H always functions well for designed proteins, indicating that yeast expression may be good.

### Implementation

The software workflow runs on a Linux server (Ubuntu 16.04 LTS). The main packages used for the implementation are TMHMM 2.0, ProPAS, RaptorX-Propert, NCBI BLAST 2.6.0+, and Perl 5.22.1 together with Bioperl modules. The responsive user interface is implemented using HTML 5 and JavaScript. To use the PSS web server, only a computer with an internet connection and a modern browser (e.g., the latest versions of Chrome, Safari, or Firefox) is required. A valid email address is needed for receiving the design results. Adobe Reader and Notepad++ may be used to view the designs. The PSS server may be accessed via http://pss.sjtu.edu.cn.

## EVALUATION AND DISCUSSION

The QTY design was originally conducted with command-line tools written in the Perl programming language, which is executable on a variety of operating systems with a Perl interpreter installed. Because many users prefer a graphic interface, the server was built with much broader group of users in mind. Generally, it comprises 2 major parts, namely the Simple-Design module and the Library-Design module which share the same input requirements (Fig. 1). For the simple-design module, users are only required to indicate the region(s) that need to be changed, and PSS will perform a thorough QTY substitution in the selected region(s). Comparatively, the Library-Design module will perform a partial substitution in order to maintain the best possible balance between solubility of the protein and its structural integrity. The Library-Design output may be used to direct library synthesis followed by yeast-two-hybrid screening (Y2H).

### Convenience evaluation

With the PSS server, one can perform designs by simply typing/pasting the sequences of target proteins. A user can also perform designs using only the UniProt number of a target protein. There are 3 different ways to input secondary structure data including the 3-state format SS3(13) (Supplementary File 1). PSS allows customized designs via input secondary structure data and/or regions that the user wants to modify. With proper input, PSS can perform the design rapidly. Finally, it will produce a detailed report for the design, in which differences between the QTY variant (MT) and the wild-type (WT) proteins can be easily checked. File(s) containing designed sequence(s) in plain text format are also provided for gene design and related DNA synthesis.

### Performance and potential

The development of PSS began in 2013 and the main part was completed in 2016. Post 2016, many QTY designs were conducted using the server. Experiments aimed to test the design were also performed, including the investigations reported in our previous study(6). Currently, we have completed all designs on human GPCRs, including related statistical analyses (Supplementary File 4). A distribution map containing these 825 human GPCRs and their QTY variants is shown (Fig. 6). Apparently, QTY-designed GPCRs are distinctly separated from their corresponding origins by hydrophobicity (H_Y_), although the α-helix ratios among the whole protein secondary structure (R_H_) vary from 0.02 to 0.78. Moreover, most thin lines are nearly vertical, strongly suggesting that only minor changes in the secondary structure were produced in the QTY design. The figure also shows that the R_H_ values of most human GPCRs range between 0.5 and 0.7, where the best water solubility improvement was predicted. This suggests that PSS based on the QTY method possesses an excellent capability for solubilizing GPCRs.

**Figure 6.**
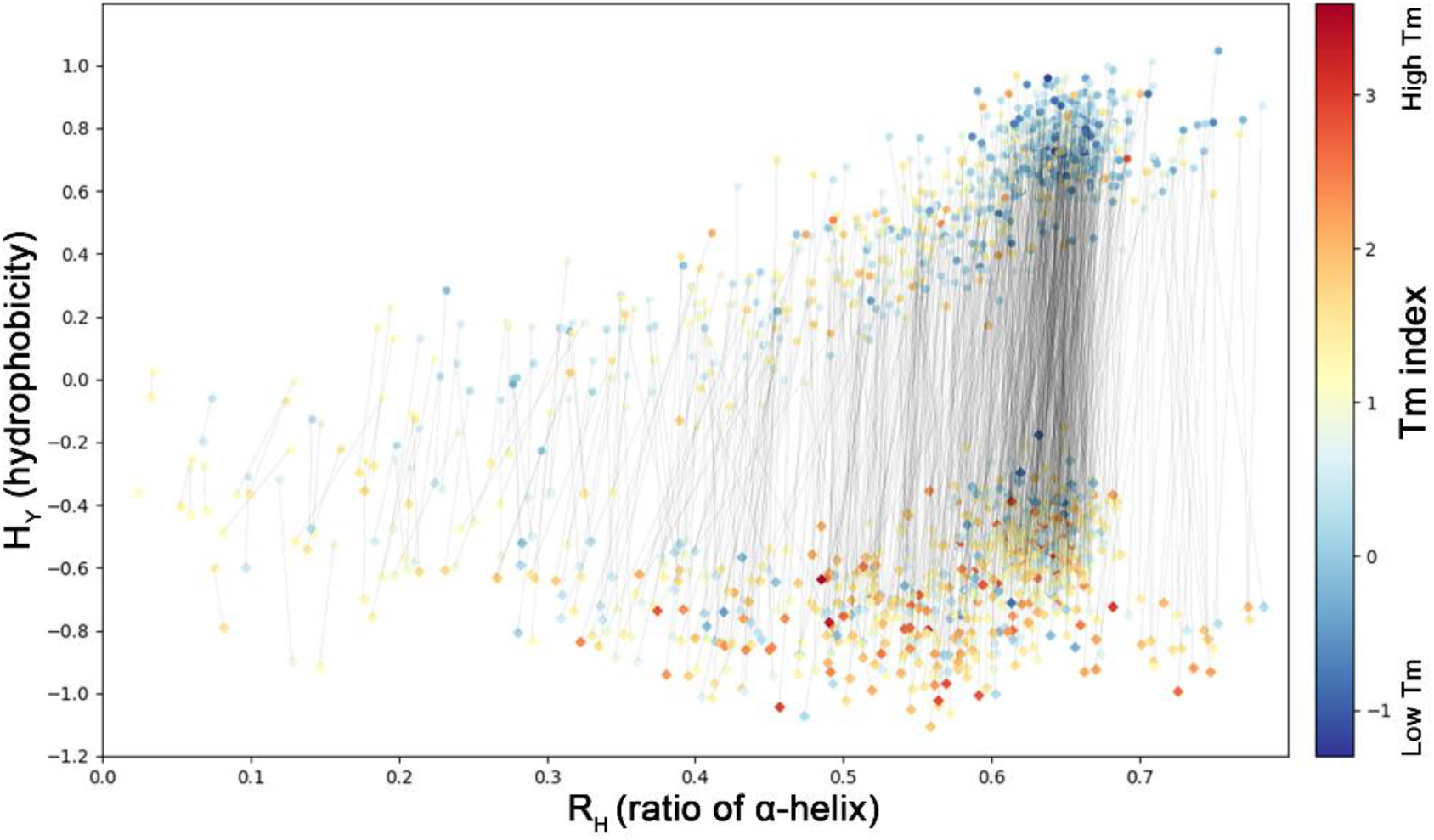
Global R_H_-H_Y_ distribution of 825 human GPCRs and their variants. The R_H_ value indicates the abundance of α-helical regions in a protein. It was calculated via dividing summarized lengths of all α-helices (including non-transmembrane helices) by protein lengths. The hydrophobicity (H_Y_) values were calculated using a standalone software ProPAS and then used for evaluating water solubility of a protein. The Tm index, shown using a color gradient, was calculated using a sequence-based method, which qualitatively represents the stability of a protein. The original GPCRs are denoted by a circle, and the QTY designed variants by a diamond. The thin black line is used to show the corresponding relationship between the original protein and its variant. The line slope represents the change rate of α-helical ratio which can partially reflect the effect of the QTY design on protein secondary structure.

The Tm index(19) of QTY designed proteins were mostly higher than their coordinates, indicating that better stabilities QTY proteins were more stable, which is highly consistent with our previous observation(6). Designing proteins with enhanced thermo-stability is a main focus of protein engineering owing to its theoretical and practical significance(20,21). Therefore, QTY-based PSS may also provide a general strategy for stabilizing proteins, which is a critical requirement(20).

PSS was designed for modifying proteins containing α-helices. Therefore, besides GPCRs, any protein with a hydrophobic α-helical secondary structure may be suitable for PSS. There are many such important proteins such as adiponectin receptors, claudin and tetraspanin. Enzymes with hydrophobic α-helices such as cellulase CelDZ1, β-carotene 15,15’-dioxygenase, and endo-β-*N*-acetylglucosaminidase(22), are also within the application scope of PSS. Although the examples listed are all TM proteins, the scope of application is beyond TM proteins. This because α-helices are also abundant in non-membrane proteins. Theoretically, these water insoluble proteins containing hydrophobic α-helical structures may also be solubilized using PSS. For this, a user needs to manually input secondary structure data as described in the procedure section.

### Limitations

The server was developed based on theories concerning protein α-helical secondary structure, and therefore, may only be used for solubilizing proteins with hydrophobic α-helical structure(s). Other types of proteins such as those with β-barrels are not suitable for PSS. Another limitation was computing capability issue, where only a single sequence per submission was allowed. In addition, each sequence could not contain more than 8,000 amino acids. The Library Design is applicable only for proteins whose variants can be screened using high-throughput methods such as Y2H method. Currently, the parameters of Library Design have only been optimized with GPCRs as the target protein and the library capacity is set at 2 million.

## SUPPLEMENTARY DATA

Supplemental File 1: Examples of submission data.

Supplemental File 2: A simple design report with annotations

Supplemental File 3: A library design result with annotations.

Supplemental File 4: Designs of 825 human GPCRs.

Supplemental File 5: GPCRs with protein or petide ligand.

Supplemental File 6: Detail procedures for both the simple and library design.

### ACKNOWLEDGMENT

Authors would like to thank Rui Qing (MIT) Jun Ni (SJTU) for their help in doing the test and evaluation of PSS server.

## FUNDING

This work was supported by Shanghai Rising-Star Program (18QA1401900), and NSFC (31870088, 31570101).

## AUTHOR CONTRIBUTIONS

F.T. built the web server, drafted and revised the manuscript. H.T. wrote and revised the manuscript. S.Z. provided original idea of QTY, and contributed a lot of advices in improving the web server and writing the paper. P.X. provided research resources, supervised the project, wrote and revised the manuscript.

## COMPETING FINANCIAL INTERESTS

F.T. is a full-time employee of Shanghai Jiao Tong University, and the University licenses all the programs related to this server. However, all programs and servers described here are free for academic use.

## REFERENCES

1. Krogh, A., Larsson, B., von Heijne, G. and Sonnhammer, E.L. (2001) Predicting transmembrane protein topology with a hidden Markov model: application to complete genomes. J. Mol. Biol., 305, 567–580.

2. Pandey, A., Shin, K., Patterson, R.E., Liu, X.Q. and Rainey, J.K. (2016) Current strategies for protein production and purification enabling membrane protein structural biology. Biochem. Cell Biol., 94, 507–527.

3. Almen, M.S., Nordstrom, K.J., Fredriksson, R. and Schioth, H.B. (2009) Mapping the human membrane proteome: a majority of the human membrane proteins can be classified according to function and evolutionary origin. BMC Biol., 7, 50.

4. Jacoby, E., Bouhelal, R., Gerspacher, M. and Seuwen, K. (2006) The 7 TM G-protein-coupled receptor target family. ChemMedChem, 1, 761–782.

5. Latorraca, N.R., Venkatakrishnan, A.J. and Dror, R.O. (2017) GPCR dynamics: structures in motion. Chem. Rev., 117, 139–155.

6. Zhang, S., Tao, F., Qing, R., Tang, H., Skuhersky, M., Corin, K., Tegler, L., Wassie, A., Wassie, B., Kwon, Y. et al. (2018) QTY code enables design of detergent-free chemokine receptors that retain ligand-binding activities. Proc. Natl. Acad. Sci. USA, 201811031.

7. Moller, S., Croning, M.D. and Apweiler, R. (2001) Evaluation of methods for the prediction of membrane spanning regions. Bioinformatics, 17, 646–653.

8. Sonnhammer, E.L., von Heijne, G. and Krogh, A. (1998) A hidden Markov model for predicting transmembrane helices in protein sequences. Proc. Int. Conf. Intell. Syst. Mol. Biol., 6, 175–182.

9. Wu, S. and Zhu, Y. (2012) ProPAS: standalone software to analyze protein properties. Bioinformation, 8, 167–169.

10. Gasteiger, E., Hoogland, C., Gattiker, A., Duvaud, S., Wilkins, M.R., Appel, R.D. and Bairoch, A. (2005) Protein identification and analysis tools on the ExPASy server. Humana Press Totowa, NJ.

11. Crabtree, J., Agrawal, S., Mahurkar, A., Myers, G.S., Rasko, D.A. and White, O. (2014) Circleator: flexible circular visualization of genome-associated data with BioPerl and SVG. Bioinformatics, 30, 3125–3127.

12. Omasits, U., Ahrens, C.H., Muller, S. and Wollscheid, B. (2014) Protter: interactive protein feature visualization and integration with experimental proteomic data. Bioinformatics, 30, 884–886.

13. Wang, S., Li, W., Liu, S. and Xu, J. (2016) RaptorX-Property: a web server for protein structure property prediction. Nucleic Acids Res., 44, W430–435.

14. Yang, Y., Gao, J., Wang, J., Heffernan, R., Hanson, J., Paliwal, K. and Zhou, Y. (2018) Sixty-five years of the long march in protein secondary structure prediction: the final stretch? Brief. Bioinformatics, 19, 482–494.

15. Kallberg, M., Wang, H.P., Wang, S., Peng, J., Wang, Z.Y., Lu, H. and Xu, J.B. (2012) Template-based protein structure modeling using the RaptorX web server. Nat. Protoc., 7, 1511–1522.

16. Yang, M., Wu, Z. and Fields, S. (1995) Protein-peptide interactions analyzed with the yeast two-hybrid system. Nucleic Acids Res., 23, 1152–1156.

17. Grote, A., Hiller, K., Scheer, M., Munch, R., Nortemann, B., Hempel, D.C. and Jahn, D. (2005) JCat: a novel tool to adapt codon usage of a target gene to its potential expression host. Nucleic Acids Res., 33, W526–531.

18. Puigbo, P., Guzman, E., Romeu, A. and Garcia-Vallve, S. (2007) OPTIMIZER: a web server for optimizing the codon usage of DNA sequences. Nucleic Acids Res., 35, W126–131.

19. Ku, T., Lu, P., Chan, C., Wang, T., Lai, S., Lyu, P. and Hsiao, N. (2009) Predicting melting temperature directly from protein sequences. Comput. Biol. Chem., 33, 445–450.

20. Li, Y., Zhang, J., Tai, D., Middaugh, C.R., Zhang, Y. and Fang, J. (2012) PROTS: a fragment based protein thermo-stability potential. Proteins, 80, 81–92.

21. Gromiha, M.M., Pathak, M.C., Saraboji, K., Ortlund, E.A. and Gaucher, E.A. (2013) Hydrophobic environment is a key factor for the stability of thermophilic proteins. Proteins, 81, 715–721.

22. Garrido, D., Nwosu, C., Ruiz-Moyano, S., Aldredge, D., German, J.B., Lebrilla, C.B. and Mills, D.A. (2012) Endo-beta-*N*-acetylglucosaminidases from infant gut-associated bifidobacteria release complex *N*-glycans from human milk glycoproteins. Mol. Cell. Proteomics, 11, 775–785.

