## Supplemental File 2 for "PSS: An enabling QTY server for designing water-soluble α-helical transmembrane proteins"

### REPORT OF QTY/NTY DESIGN

#### 1. Job information:

Job/Protein Name: P51681

User:

Designing code: QTY

Length of protein sequence: 352

[illegible]r of modified TM regions: 7

This is the name input by users. Here I used the Uniprot number as the protein name. This can be anything you want. But, one thing need to be aware. If you use the same name and the same email for different submissions. Only the last job will be kept for one week. All the submissions will be deleted after one week.

Means QTY was used in this design, it can also be NTY.

Means the original protein

#### 2. Comparison of general characteristics:

This is for estimating the overall hydrophobicity of a protein. Kyte-Doolittle scale is used as the hydrophobic index.

| Type | pI | MW (kDa) | Hydrophobicity | Mutation rate (TM, %) | Mutation rate (%) |
|---|---|---|---|---|---|
| WT | 9.20 | 40.5241 | 0.5466 | / | / |
| MT | 9.07 | 41.0524 | -0.9218 | 55.76 | 26.14 |

#### 3. Comparison of TM prediction:

Means the QTY designed protein.

Means the mutation rate of trans-membrane regions. It was calculated by dividing the number of changed AA residues by the total number of residues in transmembrane regions

It was calculated by dividing the number of changed AA residues by the length of the protein sequence

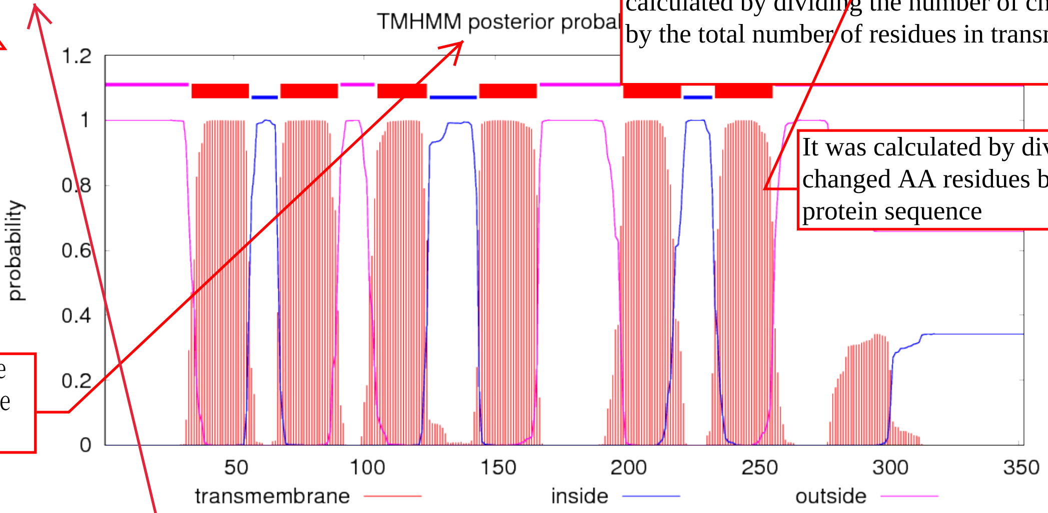


Was calculated using the wild type sequence as the input

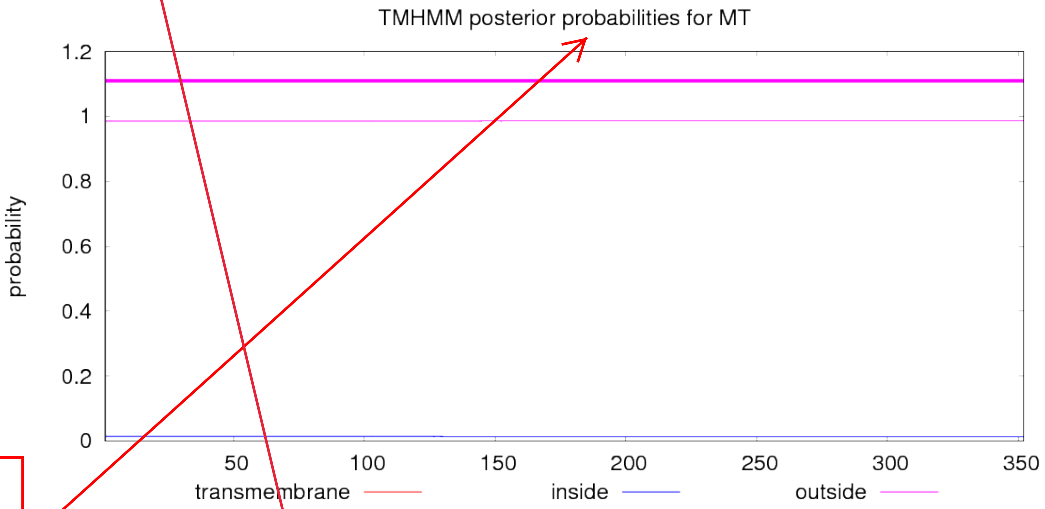


Was calculated using the variant sequence as the input.

This is done with the help of TMHMM 2.0. Every peak corresponds to a TM region. If there is no peak showing in the figure, that means all the TM regions have been removed from the protein.

#### 4. Comparison of sequences:

Means identical

Means different

```
  1                                                                60
WT  MDYQVSSPIYDINYYTSEPCQKINVKQIAARLLPPLYSLVFIFGFVGNMLVILILINCKR
    |||||||||||||||||||||||||||||||**||*||*****|**|||*******||||
MT  MDYQVSSPIYDINYYTSEPCQKINVKQIAARQQPPQYSQTYTYGYTGNMQTTQTQTNCKR

 61                                                               120
WT  LKSMTDIYLLNLAISDLFFLLTVPFWAHYAAAQWDFGNTMCQLLTGLYFIGFFSGIFFII
    ||||||||**|*|*||*****|*|*|||||||||||||||||**||*|**|**||*****
MT  LKSMTDIYQQNQATSDQYYQQTTPYWAHYAAAQWDFGNTMCQQQTGQYYTGYYSGTYYTT

121                                                               180
WT  LLTIDRYLAVVHAVFALKARTVTFGVVTSVITWVVAVFASLPGIIFTRSQKEGLHYTCSS
    **|*|||||||||||||||||*|*|**||**||**|**||*||***||||||||||||||
MT  QQTTDRYLAVVHAVFALKARTTTYGTTTSTTTWTTATYASQPGTTYTRSQKEGLHYTCSS

181                                                               240
WT  HFPYSQYQFWKNFQTLKIVILGLVLPLLVMVICYSGILKTLLRCRNEKKRHRAVRLIFTI
    ||||||||||||||||||***|***|***|**||||**|||||||||||||||||***|*
MT  HFPYSQYQFWKNFQTLKITTQGQTQPQQTMTTCYSGTQKTLLRCRNEKKRHRAVRQTYTT

241                                                               300
WT  MIVYFLFWAPYNIVLLLNTFQEFFGLNNCSSSNRLDQAMQVTETLGMTHCCINPIIYAFV
    |**|***|||||*****||*||||||||||||||||||||*|||*||||||*||**||**
MT  MTTYYQYWAPYNTTQQQNTYQEFFGLNNCSSSNRLDQAMQTTETQGMTHCCTNPTTYAYT

301                                                               360
WT  GEKFRNYLLVFFQKHIAKRFCKCCSIFQQEAPERASSVYTRSTGEQEISVGL
    ||||||||||||||||||||||||||||||||||||||||||||||||||||
MT  GEKFRNYLLVFFQKHIAKRFCKCCSIFQQEAPERASSVYTRSTGEQEISVGL
```

#### 5. Comparison of Protter prediction:

This is completed by using the Protter sever. The TM region prediction and sub-cellular localization are all done by the server. It can clearly show that if there are still TM regions in a protein and if the protein is still in membrane.

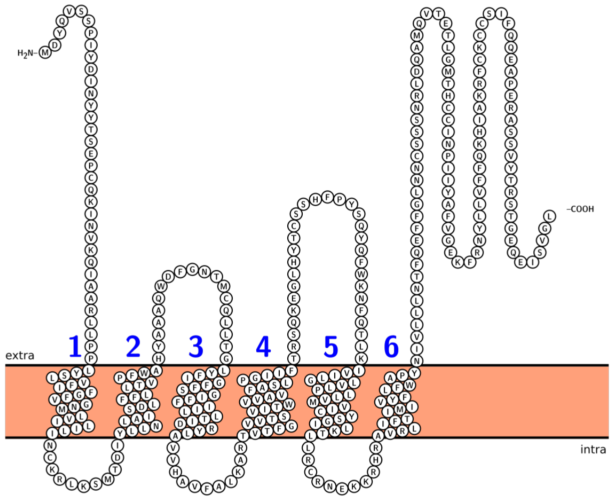


WT

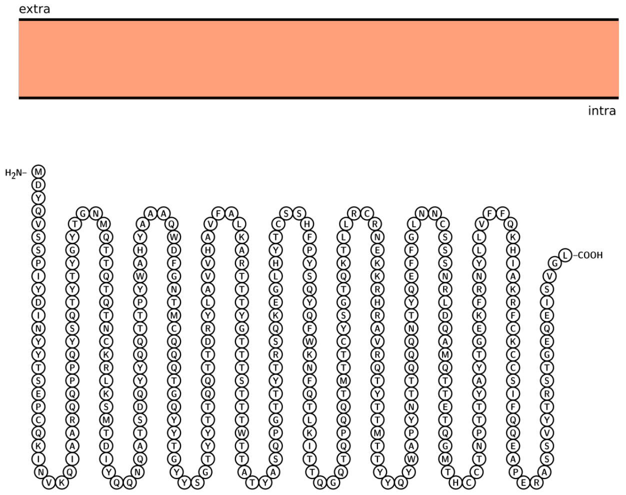


MT

#### 6. Detailed comparison of TM regions:

Indicate the first trans-membrane helix

### TM1

**AA sequence comparison:**

```
TM1-wt:  LLPPLYSLVFIFGFVGNMLVILILINCK
         **||*||*****|**|||*******|||
TM1-mt:  QQPPQYSQTYTYGYTGNMQTTQTQTNCK
```

**Alpha-helix prediction comparison:**

The prediction was done with RaptorX

```
TM1-wt:  HHHHHHHHHHHHHHHHHHHHHHHHHHHC
         |||||||||||||||||||||||||||*
TM1-mt:  HHHHHHHHHHHHHHHHHHHHHHHHHHHH
```

"H" means helix

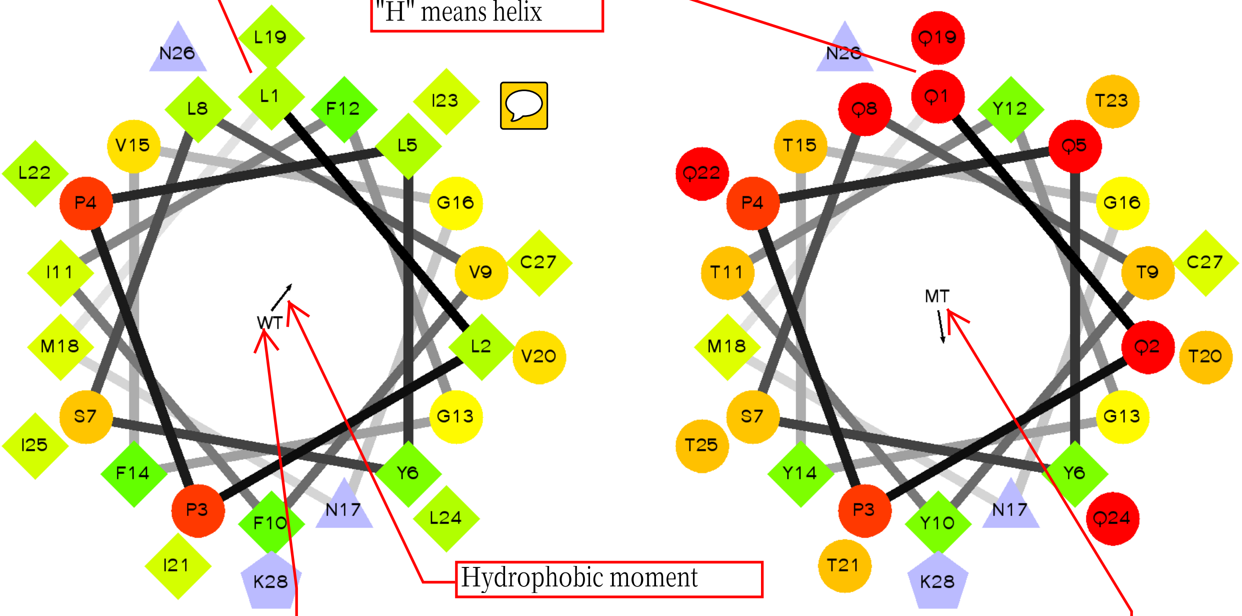


Wheel of the helix of wild type trans-membrane region

Wheel of the helix of QTY designed trans-membrane region

### TM2

**AA sequence comparison:**

```
TM2-wt:  LNLAISDLFFLLTVPFWAHYA
         *|*|*||*****|*|*|||||
TM2-mt:  QNQATSDQYYQQTTPYWAHYA
```

**Alpha-helix prediction comparison:**

"C" means coil

```
TM2-wt:  HHHHHHHHHHHCCCHHHHHHH
         |||||||||||||||||||||
TM2-mt:  HHHHHHHHHHHCCCHHHHHHH
```

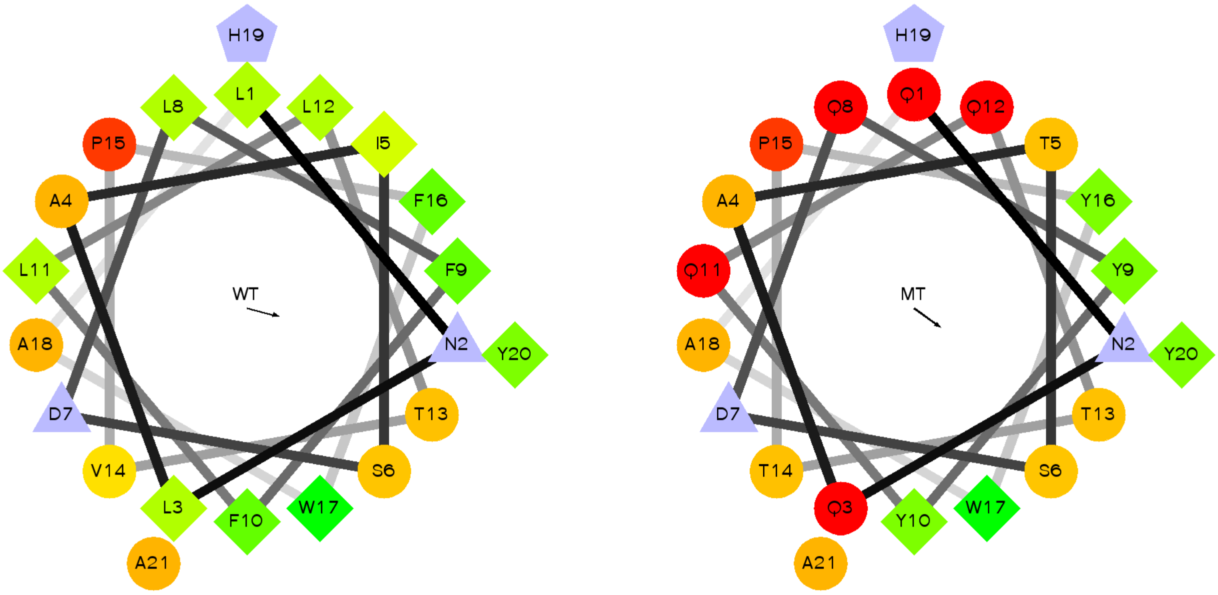

# TM3

#### AA sequence comparison:

```
TM3-wt:  LTGLYFIGFFSGIFFIILLTID
         *||*|**|**||*******|*|
TM3-mt:  QTGQYYTGYYSGTYYTTQQTTD
```

#### Alpha-helix prediction comparison:

```
TM3-wt:  HHHHHHHHHHHHHHHHHHHHHH
         ||||||||||||||||||||||
TM3-mt:  HHHHHHHHHHHHHHHHHHHHHH
```

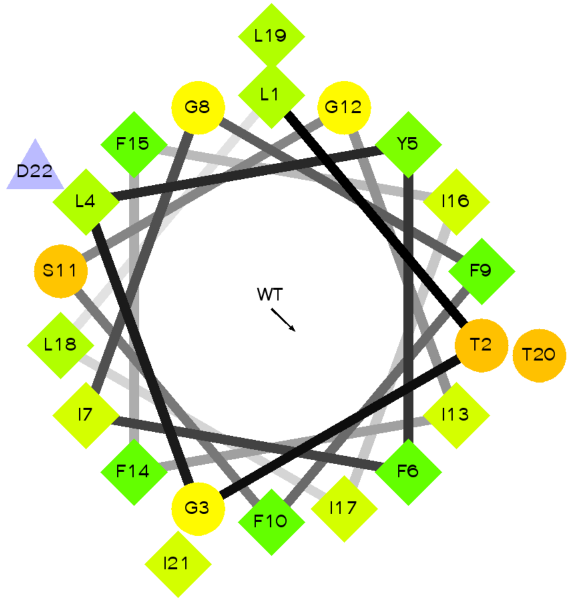


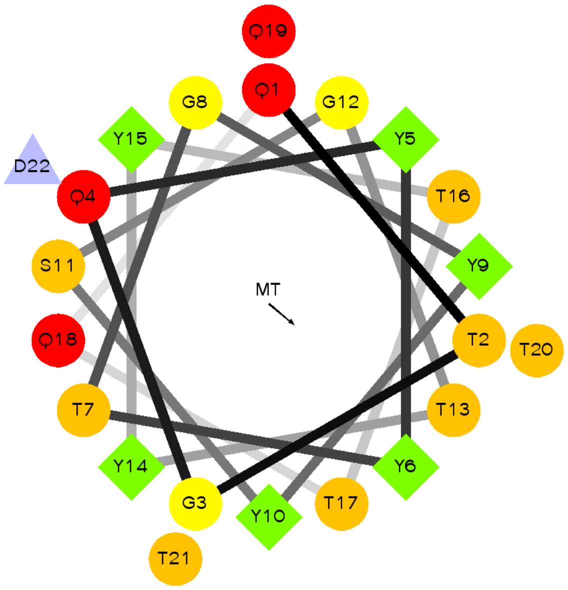


# TM4

#### AA sequence comparison:

```
TM4-wt:  TFGVVTSVITWVVAVFASLPGIIFT
         |*|**||**||**|**||*||***|
TM4-mt:  TYGTTTSTTTWTTATYASQPGTTYT
```

"E" means belta-sheet

#### Alpha-helix prediction comparison:

```
TM4-wt:  HHHHHHHHHHHHHHHHHHCCCEEEE
         |||||||||||||||||*|||||||
TM4-mt:  HHHHHHHHHHHHHHHHHCCCCEEEE
```

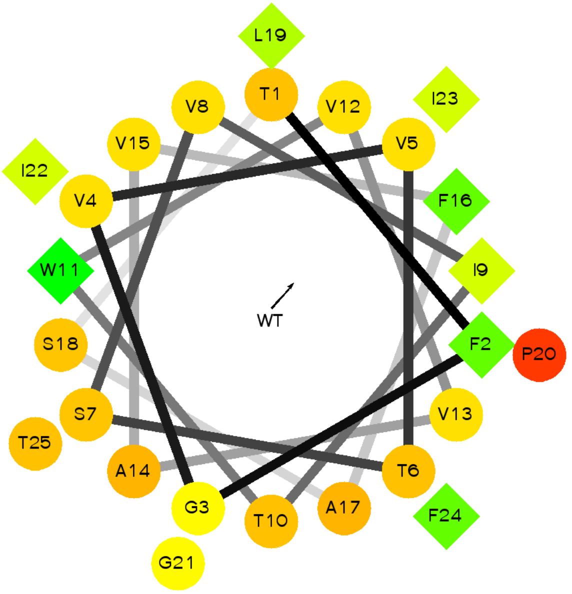


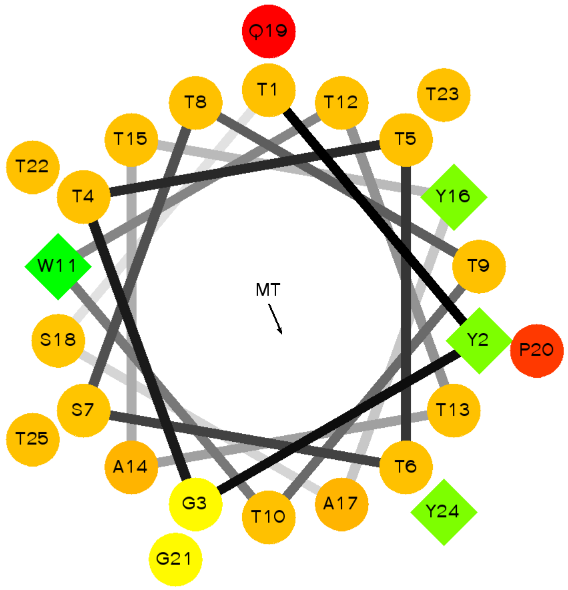

## TM5

##### AA sequence comparison:

```
TM5-wt:  ILGLVLPLLVMVICYSGILK
         **|***|***|**||||**|
TM5-mt:  TQGQTQPQQTMTTCYSGTQK
```

##### Alpha-helix prediction comparison:

```
TM5-wt:  HHHHHHHHHHHHHHHHHHHH
         ||||||||||||||||||||
TM5-mt:  HHHHHHHHHHHHHHHHHHHH
```

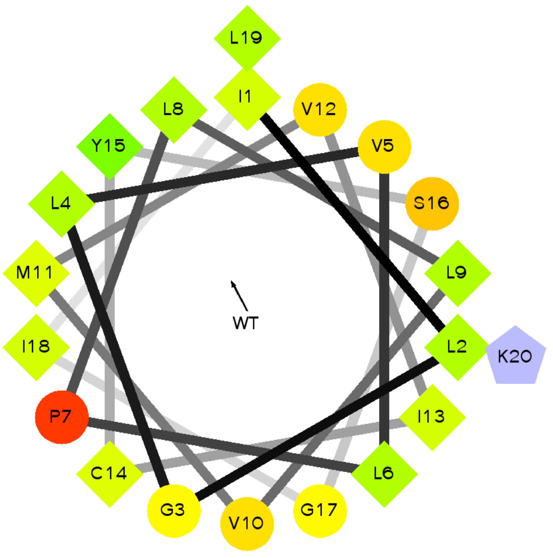


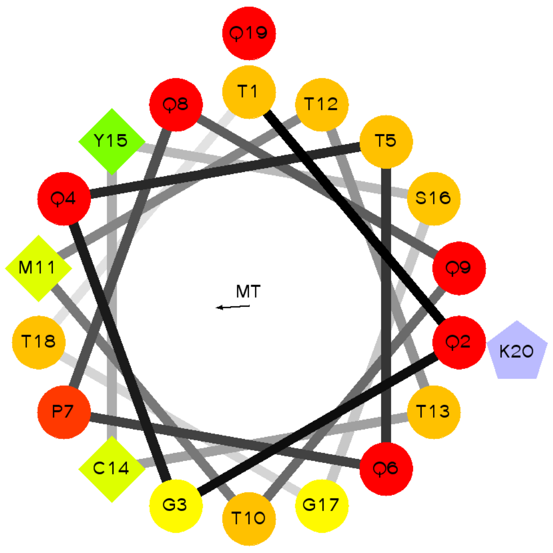


## TM6

##### AA sequence comparison:

```
TM6-wt:  IFTIMIVYFLFWAPYNIVLLLNTFQ
         **|*|**|***|||||*****||*|
TM6-mt:  TYTTMTTYYQYWAPYNTTQQQNTYQ
```

##### Alpha-helix prediction comparison:

```
TM6-wt:  HHHHHHHHHHHCCHHHHHHHHHHHH
         |||||||||||||||||||||||||
TM6-mt:  HHHHHHHHHHHCCHHHHHHHHHHHH
```

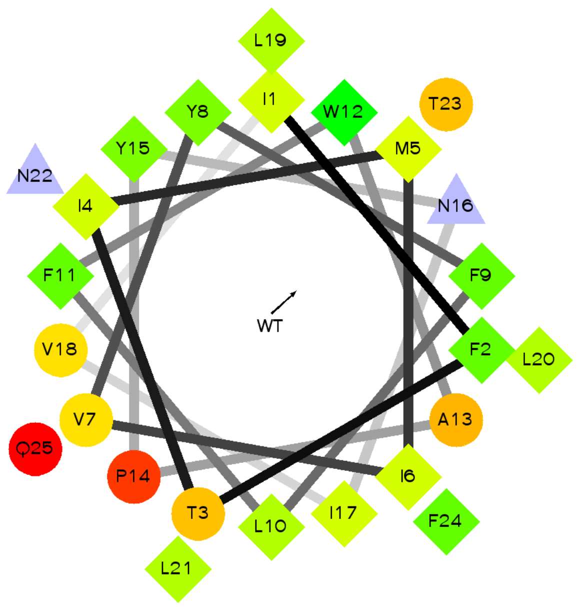


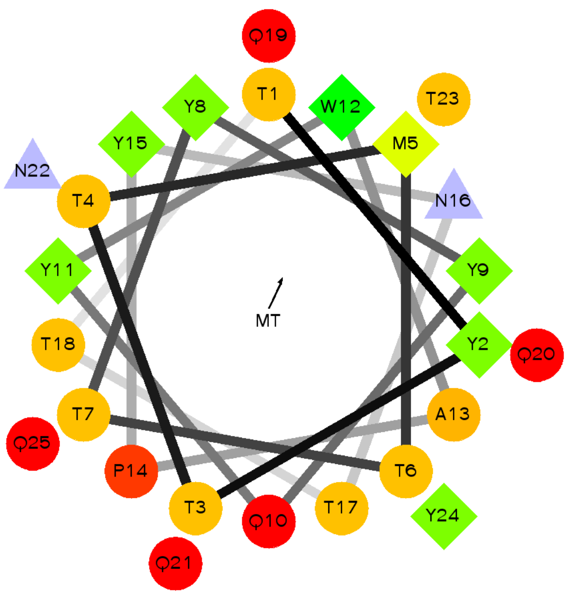

# TM7

#### AA sequence comparison:

```
TM7-wt:  MQVTETLGMTHCCINPIIYAFVGE
         ||*|||*||||||*||**||**||
TM7-mt:  MQTTETQGMTHCCTNPTTYAYTGE
```

#### Alpha-helix prediction comparison:

```
TM7-wt:  HHHHHHHHHHHHHHHHHHHHHHCH
         |||||||||||****||||||*|*
TM7-mt:  HHHHHHHHHHHCCCCHHHHHHCCC
```

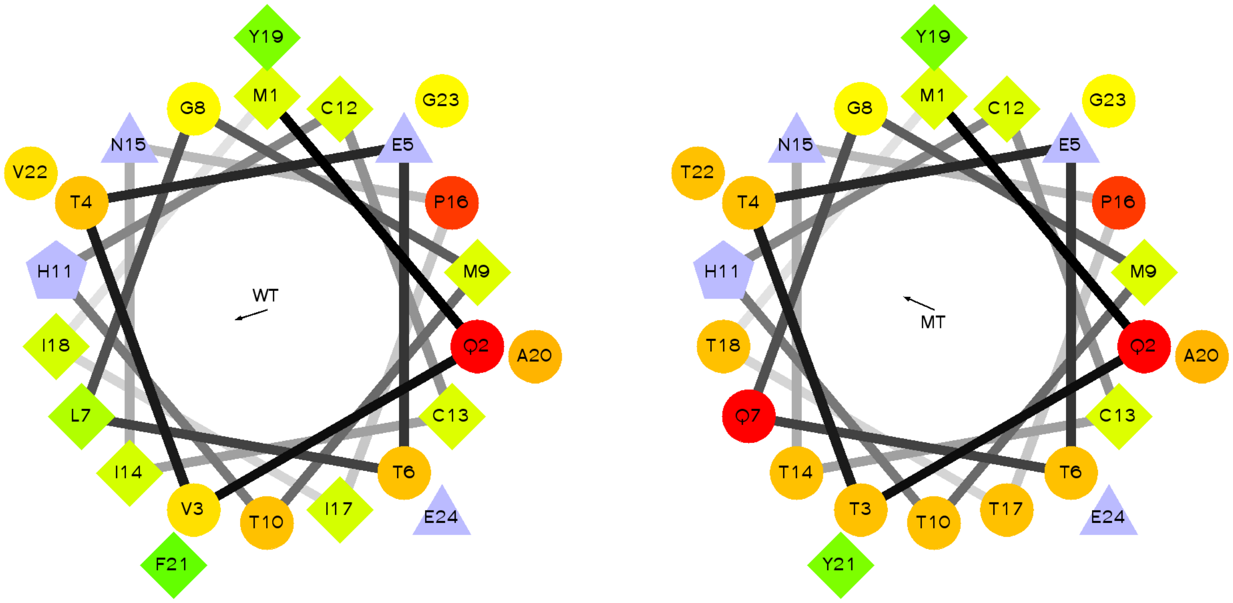
