## Supplemental File 3 for "PSS: An enabling QTY server for designing water-soluble α-helical transmembrane proteins"

This is the ".out.txt" file which can serve for the printing.

Notepad++ is strongly suggested for opening the results.

The name contains information of the job name and design code.

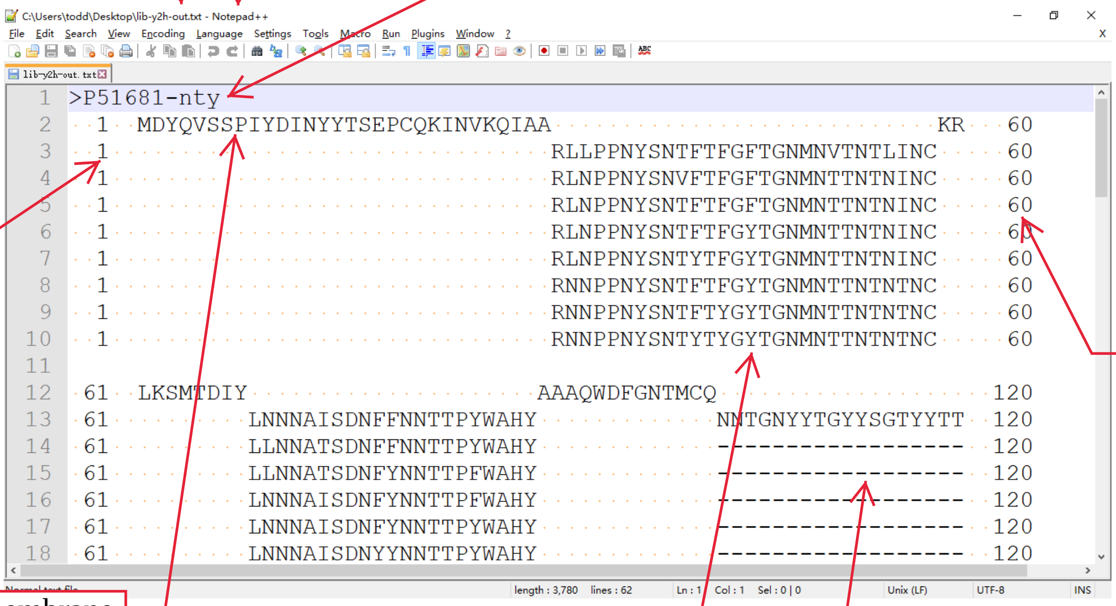


The position of amino acid

The position of amino acid

The non-transmembrane region. These regions are identical to original protein.

This is of the TM region. There are 8 different variants for this TM region.

As for this TM region, there are only one variant which can be designed. This is because changing more AAs will lead to a predicted structure change.

This is a full view of the whole result of the library design. It is easy to find that there are totally 7 TM regions and 8 non-TM regions.

```
>P51681-nty
  1  MDYQVSSPIYDINYYTSEPCQKINVKQIAA                            KR    60
  1                                RLLPPNYSNTFTFGFTGNMNVTNTLINC      60
  1                                RLNPPNYSNVFTFGFTGNMNTTNTNINC      60
  1                                RLNPPNYSNTFTFGFTGNMNTTNTNINC      60
  1                                RLNPPNYSNTFTFGYTGNMNTTNTNINC      60
  1                                RLNPPNYSNTYTFGYTGNMNTTNTNINC      60
  1                                RNNPPNYSNTFTFGYTGNMNTTNTNTNC      60
  1                                RNNPPNYSNTFTYGYTGNMNTTNTNTNC      60
  1                                RNNPPNYSNTYTYGYTGNMNTTNTNTNC      60

 61  LKSMTDIY                     AAAQWDFGNTMCQ                     120
 61          LNNNAISDNFFNNTTPYWAHY             NNTGNYYTGYYSGTYYTT  120
 61          LLNNATSDNFFNNTTPYWAHY             ------------------  120
 61          LLNNATSDNFYNNTTPFWAHY             ------------------  120
 61          LNNNAISDNFYNNTTPFWAHY             ------------------  120
 61          LNNNAISDNFYNNTTPYWAHY             ------------------  120
 61          LNNNAISDNYYNNTTPYWAHY             ------------------  120
 61          LNNNATSDNYYNNTTPYWAHY             ------------------  120
 61          NNNNATSDNYYNNTTPYWAHY             ------------------  120

121      DRYLAVVHAVFALKART                        TRSQKEGLHYTCSS  180
121  NNTT                 TTYGTTTSTTTWTTATYASNPGTTY                180
121  ----                 -------------------------                180
121  ----                 -------------------------                180
121  ----                 -------------------------                180
121  ----                 -------------------------                180
121  ----                 -------------------------                180
121  ----                 -------------------------                180
121  ----                 -------------------------                180

181  HFPYSQYQFWKNFQTLKI                    KTLLRCRNEKKRHRAVR        240
181                    VINGNVNPNNVMTTCYSGTL                 NTFTT  240
181                    VTNGNVNPNNTMVTCYSGIN                 NTFTT  240
181                    TINGNVNPNNTMVTCYSGTL                 NTFTT  240
181                    VTNGNVNPNNTMTICYSGTL                 NTFTT  240
181                    VTNGNVNPNNTMTTCYSGIN                 NTYTT  240
181                    TINGNTNPNNTMTTCYSGTL                 NTYTT  240
181                    TTNGNTNPNNTMVTCYSGTN                 NTYTT  240
181                    TTNGNTNPNNTMTTCYSGTN                 NTYTT  240

241                      QEFFGLNNCSSSNRLDQ                             300
241  MTTYFNFWAPYNTVNNLNTF                 AMQVTETNGMTHCCINPIIYAFV  300
241  MTTYYNFWAPYNTVNNLNTF                 AMQVTETNGMTHCCTNPIIYAFV  300
241  MTTYYNYWAPYNTVNNLNTF                 AMQVTETLGMTHCCTNPITYAFT  300
241  MTTYYNFWAPYNTVNNNNTF                 AMQVTETNGMTHCCTNPIIYAFT  300
241  MTTYYNFWAPYNTVNNNNTF                 AMQVTETNGMTHCCTNPTIYAFT  300
241  MTTYYNYWAPYNTVNNNNTF                 AMQVTETNGMTHCCTNPITYAYT  300
241  MTTYYNYWAPYNTTNNNNTF                 AMQTTETNGMTHCCTNPTTYAYV  300
241  MTTYYNYWAPYNTTNNNNTY                 AMQTTETNGMTHCCTNPTTYAYT  300

301   EKFRNYLLVFFQKHIAKRFCKCCSIFQQEAPERASSVYTRSTGEQEISVGL  352
301  G                                                    352
301  G                                                    352
301  G                                                    352
301  G                                                    352
301  G                                                    352
301  G                                                    352
301  G                                                    352
301  G                                                    352
```

The name of this file contains "out2" . It is designed for facilitating the following sequence manipulations.

Here, we also recommend using the Notepad++ for the treatment of the sequence.

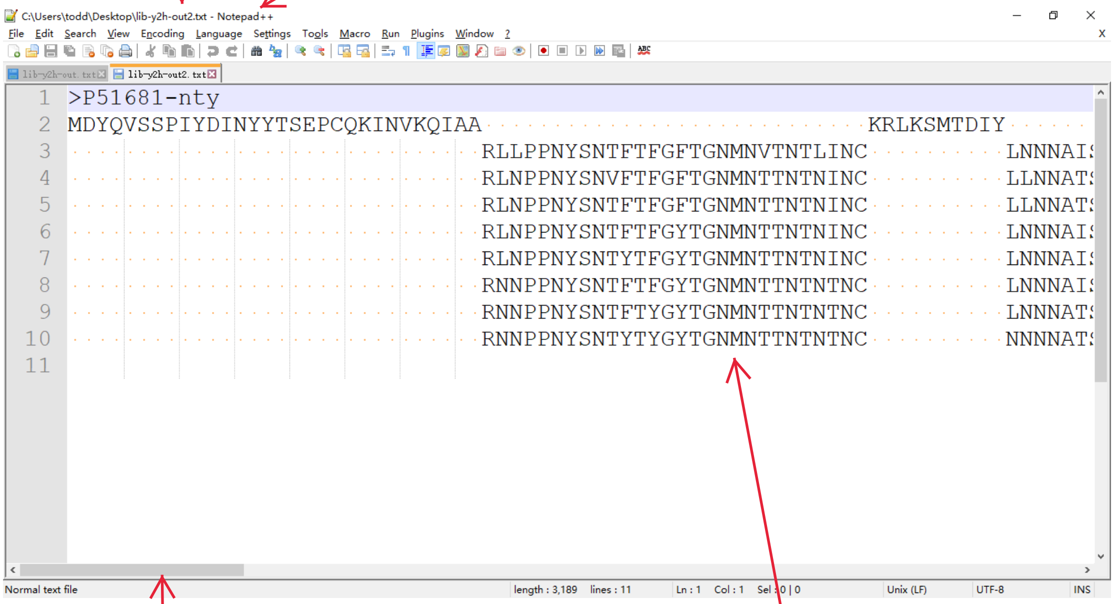


As for this file, you can scroll the bar to see the whole sequence.

The 8 variants of the TM region.
