## Supplemental File 6 for "PSS: An enabling QTY server for designing water-soluble α-helical transmembrane proteins"

**PROCEDURE**

**Preparation of required information ● TIMING ~5 min**

**1 |** This step varies depending on a user’s situation in real time. In general, 3 types of information are needed:

**(A) Amino acid sequence of target protein**

Protein sequences to be submitted may be downloaded or copied from any source, such as NCBI. The sequence may be in FASTA format or in string format. Any notation characters embedded in the sequence will be removed automatically by the server. For convenience, the UniProt number (protein record ID) may be used as input. In case the UniProt number is input, the server will retrieve the UniProt database and acquire the canonical sequence. A UniProt number can be obtained by performing a simple search on the UniProt website (www.uniprot.org).

**(B) α-Helical TM region information**

This section is optional because we have integrated TM region prediction software and TM region retrieving programs in PSS. However, accurate TM region information is critical for an effective QTY design. If a user obtains accurate TM region information from crystal structure data or from another method, we recommend that the data be supplied. PSS accepts 2 types of TM region information data, SS3 format and start-end position format. The SS3 format string can be obtained from many sources such as RaptorX prediction. This format uses 3 letters to indicate the secondary structure of each position, where “H” stands for helix, “C” for coli, and “E” for unknown structure. The start-end position format simply provides the start and end positions of each TM fragments (SI-1).

**▲CRITICAL STEP** For both formats provided above, all regions labeled as ‘helix’ or ‘H’ will be recognized as TM helices and will be exposed to QTY substitution. The α-helix is also known to exist in non-TM proteins. Therefore, in data containing secondary structure of a protein with non-TM α-helices, the section ‘Specify region(s) to do design’ must be used to accurately indicating TM regions in order to avoid changing the non-TM helices. The secondary structure data may also be edited by only labeling the TM helices with ‘helix’ or ‘H’.

**(C) Desired mutant region(s)**

This section is also optional, as it is only meant for partial modification of a protein without affecting some of the α-helices. An input box in the webpage was designed to enable start-end position pairs to be entered in order to indicate the regions requiring QTY modification. Thus, PSS will only change α-helices in the indicated regions and leave the other parts as they were in wild type.

**Submitting a job ● TIMING 5 min**

**2 |** Go to the PSS homepage at http://pss.sjtu.edu.cn/.

**3 |** Select ‘Design’ from the menu at the left of the page.

**4 |** Select between ‘Simple Design’ and ‘Library Design’.

**▲CRITICAL STEP** By default, the ‘Simple Design’ is selected. This is the classical QTY design and can be completed in minutes. The ‘Library Design’ is only applicable for proteins that can be screened via high-throughput methods.

**5 |** Supply a job name for identification of different submissions, and an e-mail address in order to receive notifications and design results when the design has been completed.

**▲CRITICAL STEP** PSS does not require a user to register before submitting a job. Therefore, it is important to provide a correct e-mail address. Otherwise, the results will not be received. The job name is used for identifying each job. It must be ensured that different job names are used for different submissions. Otherwise, only the result of the last submission will be received.

**6 |** In the ‘Code Selection’ section, the user need to select a code. By default, the QTY code is selected. An alternative code, NTY, which is considered less effective in making the protein water-soluble but more effective in retaining the native protein structure, is also provided for the design.

**7 |** The ‘Amino acid sequence’ contains 2 text fields. The upper field is used to enter the UniProt number which can be used to obtain the corresponding sequence from the UniProt database. The lower text field, which is disabled by default, is for typing/pasting a sequence. The checkbox to the left of ‘UniProt number’, must be unchecked before typing/pasting a sequence in the text field.

**8 |** To input TM region information (the secondary structure data), 3 ways are provided. Once the UniProt number is provided, PSS will retrieve TM information from the UniProt database by default. The radio button to the right of ‘Structure data in SS3 format’ can be clicked to activate the text field for typing/pasting SS3 format string. By clicking the radio button after ‘Type/paste structure data as example’ TM data of the start-end positions may be entered.

**9 |** The section ‘Specify region(s) to do design’ is initially disabled. Usually it can be ignored. In case partial modification of the TM regions are wanted, the checkbox must be selected to activate the text field to enter the start and end positions of the desired fragment(s).

**10 |** Press the ‘Submit’ button to queue the job on PSS. A successful submission will generate a popup notification stating, ‘Your job has been successfully submitted! The result will be sent to your email address shortly’.

**▲CRITICAL STEP** Upon submission, the data entered in the form will be validated, and the user will be notified of any errors that need to be corrected in a popup message box. It must be noted that only a limited number of pending jobs are allowed for one user (as identified by their IP address used at submission) in order to maintain sufficient server capacity to serve all users. Specifically, each user may not have more than 20 Simple Design jobs and 5 Library Design pending jobs pending at any point in time. There is no API interface for PSS, and program-aided submission is not allowed.

**Job availability ● TIMING 2 min – 5 h**

**11 |** Currently, no job monitoring interface is provided. Therefore, submission of a job is followed by a waiting period. The design result will be sent to the provided e-mail address upon completion of the job. Typically, it takes 2 – 4 min for the Simple Design, while it takes 2 – 5 h for the Library Design. The accurate time cost depends on protein size and number of TM regions. This time cost does not include queuing time.

**12 |** Frequently check the provided email box until an email titled ‘QTY design for XXXX’ is received. The ‘XXXX’ stands for the job name entered during submission.

**Viewing design results ● TIMING 10 – 15 min**

**13 |** This step differs depending on whether a Simple Design (A) or Library Design (B) job is being submitted:

**(A) Viewing ‘Simple Design’ results**

(i) The e-mail contains 2 attachments. Firstly download them to a local file folder.

(ii) The file with a ‘.txt’ extension contains the sequence of the designed protein. It may be easily used in the subsequent gene design and synthesis.

(iii) The file in PDF format consists mainly of detailed comparisons between the QTY designed protein and the original protein. It contains 6 sections, where the first one contains basic information related to the job and the design. The second part is a table showing basic comparisons of the general characteristics of the 2 proteins. The third part comprises a figure containing TM region predictions for both the original protein and the designed variant. Whether the design will make the protein water-soluble may be easily checked. Part 4 is a comparison that indicates the difference between the two sequences at amino acid residue level. Part5 is the comparison of Protter predictions that indicates protein structure changes using a serpentine figure. Part 6 shows the difference between the two proteins, helix by helix. The secondary structure data, in SS3 format, is included to indicate significant changes in the secondary structure. Wheel figures of the helixes of both the original and designed proteins are embedded to illustrate the change in solubility and hydrophobic moment^23^.

**(B) Viewing ‘Library Design’ result**

(i) Download the attached files. The library design results should contain two files, namely “*.out.txt” and “*.out2.txt”. The ‘*’ is a wildcard.

(ii) Open the file “*.out.txt” using a text file editing software program such as Notepad++. The file contains sequences in in multiple line format that is suitable for direct printing.

(iii) Open the file “*.out2.txt” with a text file editing software program such as Notepad++. This file is for the subsequent DNA synthesis, and is convenient for copy/paste operations. This file contain sequences in one-line mode, which can be easily read using a PC. In the files, all variants of each helix are listed in different lines. Open the file in Notepad++, with word wrap unchecked, and the user may easily observe the relationship between different peptide fragments in the designed library.

**DNA design and synthesis ● TIMING 2 days – 3 weeks**

**14 |** This step is for reverse translating the protein sequence to DNA sequence. It differs depending on whether a Simple Design (A) or Library Design (B) job is being submitted:

**(A) DNA design for ‘Simple Design’**

Directly reverse translates the designed protein sequence to DNA sequence. This can be done with the assistance of JCat, the URL of which can be found on the ‘Tools’ page. The codon-usage bias of desired hosts and the subsequent cloning process should be considered here. Gene design services are available via many DNA synthesis company.

**(B) DNA design for ‘Library Design’**

(i) Copy the divided fragments from the ‘*.out2.txt’ file.

(ii) Reverse translate each fragment to DNA independently. Codon usage bias should be considered at this step.

(iii) Add overlapping sequences at the termini of the neighboring fragments.

(iv) Design primers for application of the DNA encoding whole length protein. The primers may be named as Pap1 and Pap2, respectively.

**15 |** This step is aimed at obtaining real DNA molecules and differs depending on whether a Simple Design (A) or Library Design (B) job is being submitted:

**(A) DNA synthesis for ‘Simple Design’**

Perform a de novo DNA chemical synthesis by acquiring a commercial service. Usually, this process takes 2 – 7 days depending on the size of DNA and the selected company.

**(B) DNA synthesis for ‘Library Design’**

(i) Chemically synthesize the designed DNA fragments separately.

(ii) Mix the synthesized DNA fragments together and perform a Non-standard PCR without Pap1 and Pap2. This is to join the fragments.

(iii) Perform PCR with Pap1 and Pap2 as the primers and the product of the last step as the template.

(iv) This entire process takes 2 – 4 weeks, depending on the protein size and selected company.

**▲CRITICAL STEP** Library synthesis is highly complicated and requires well trained technicians. Therefore, requesting professional service is strongly recommended. GenScript (Nanjing, China) and Ginkgo Bioworks (Boston, USA, previous Gen9), who have established the method by collaborating with us are recommended. Therefore, their service may be requested as per user’s convenience. The above stated 3-step assembly procedure is for normal-sized proteins. Assembly of large protein may require more steps.

**Screening with Y2H ● TIMING 10 days**

**16 |** Sub clone the library into Y2H vector to construct Y2H library.

**17 |** Perform yeast mating, and then screen the resulting cells with selection medium.

**18 |** Pick clones and verify them using colony PCR.

**19 |** Perform a re-mating for verification of screened transformants.

**20 |** Sequence the verified transformants to obtain the corresponding DNA sequences.

**▲CRITICAL STEP** Steps 16 – 20 can be skipped for ‘Simple Design’. Y2H is well established. Reference manuals are available elsewhere. However, Y2H is yet a highly technical procedure, especially at the library construction step. Therefore, a professional company who is experience in Y2H should be contacted for service. In our previous work, we performed all Y2H experiments using the services of NextInteraction (Berkeley, USA).

**Protein expression ● TIMING 3 days**

**21 |** Clone the obtained DNA into an expression vector, and then cultivate the resulting cells for protein expression.

**▲CRITICAL STEP** One is encouraged to try different methods for conducting protein expression experiments, as this procedure is still in its formative stage and only several hosts have been tried. In regard to QTY designed human GPCRs, we have tried *E. coli* hosts, and insect cells. Insect cells are recommended for use due to the absence of inclusion-body issues. Other host systems, such as yeast system, may also be effective as Y2H always performs well for designed proteins, which indicates that the expression is fine in yeast. As for QTY variants screened from Y2H, their genes often need to be redesigned. This may be performed as per step 14(A) and 15(A) with a target expression host cell in mind, such as the insect cell.

**22 |** The expression of the proteins must be determined, followed by their purification and characterization. The characterizing technique should be selected based on features of the protein concerned. For example, ligand binding was detected during the evaluation the QTY designed Chemokine receptors in our previous research^6^.
