## Supplemental File 1 for "PSS: An enabling QTY server for designing water-soluble α-helical transmembrane proteins"

**Data examples of Protein^QTY^/Library^QTY^ design**

**Example 1**

**Name:** CXCR4(C-X-C chemokine receptor type 4)

**Uniprot Number:** P61073

**Protein sequence:** (It is in the FASTA format. Simply copy, paste and save in a text editor software can make a file suitable for uploading)

>CXCR4

MEGISIYTSDNYTEEMGSGDYDSMKEPCFREENANFNKIFLPTIYSIIFLTGIVGNGLVI

LVMGYQKKLRSMTDKYRLHLSVADLLFVITLPFWAVDAVANWYFGNFLCKAVHVIYTVNL

YSSVLILAFISLDRYLAIVHATNSQRPRKLLAEKVVYVGVWIPALLLTIPDFIFANVSEA

DDRYICDRFYPNDLWVVVFQFQHIMVGLILPGIVILSCYCIIISKLSHSKGHQKRKALKT

TVILILAFFACWLPYYIGISIDSFILLEIIKQGCEFENTVHKWISITEALAFFHCCLNPI

LYAFLGAKFKTSAQHALTSVSRGSSLKILSKGKRGGHSSVSTESESSSFHSS

**Secondary structure data:** (It is in index format)

39,63,H

78,99,H

111,130,H

155,174,H

196,216,H

242,261,H

283,302,H

**Secondary structure data:** (It is in SS3 format)

CCCCCCCCCCCCCCCCCCCCCCCCCCCCCCCCCCCCCCHHHHHHHHHHHHHHHHHHHHHH

HHHCCCCCCCCCCCCCCHHHHHHHHHHHHHHHHHHHHHHCCCCCCCCCCCHHHHHHHHHH

HHHHHHHHHHCCCCEEEEEECCCCCCCCCCCCCCHHHHHHHHHHHHHHHHHHHHEECCCC

CCCEEEEEECCCCCCHHHHHHHHHHHHHHHHHHHHHCCCCCCCCCCCCCCCCCCCCCCCC

CHHHHHHHHHHHHHHHHHHHHCCCCCCCECCCCCCCCCCCCCHHHHHHHHHHHHHHHHHH

HHCCCCCCCCCCCCCCCCCCCCCCCCCCCCCCCCCCCCCCCCCCCCCCCCCC

**Example 2**

**Name:** CLDN3(Claudin-3)

**Uniprot Number:** O15551

**Protein sequence:** (It is in the FASTA format. Simply copy, paste and save in a text editor software can make a file suitable for uploading)

>CLDN3

MSMGLEITGTALAVLGWLGTIVCCALPMWRVSAFIGSNIITSQNIWEGLWMNCVVQSTGQ

MQCKVYDSLLALPQDLQAARALIVVAILLAAFGLLVALVGAQCTNCVQDDTAKAKITIVA

GVLFLLAALLTLVPVSWSANTIIRDFYNPVVPEAQKREMGAGLYVGWAAAALQLLGGALL

CCSCPPREKKYTATKVVYSAPRSTGPGASLGTGYDRKDYV

**Secondary structure data:** (It is in index format)

1,8,Cytoplasmic

9,29,Helical

30,80,Extracellular

81,101,Helical

102,115,Cytoplasmic

116,136,Helical

137,159,Extracellular

160,180,Helical

181,220,Cytoplasmic

**Secondary structure data:** (It is in SS3 format)

CCCCCCCCHHHHHHHHHHHHHHHHHHHHHCCCCCCCCCCCCCCCCCCCCCCCCCCCCCCC

CCCCCCCCCCCCCCCCCCCCHHHHHHHHHHHHHHHHHHHHHCCCCCCCCCCCCCCHHHHH

HHHHHHHHHHHHHHHHEEEECCCCCCCCCCCCCCCCCCCHHHHHHHHHHHHHHHHHHHHH

CCCEEEEEECCCCCCCCCCCCCCCCCCCCCCCCCCCCCCC

**Example 3**

Name CCR5

**Uniprot Number:** P51681

**Specified regions for design (It is in index format)**

30,150

160,180

181,220
